# Interleukin-17 and senescence regulate the foreign body response

**DOI:** 10.1101/583757

**Authors:** Liam Chung, David Maestas, Andriana Lebid, Ashlie Mageau, Gedge D. Rosson, Xinqun Wu, Matthew T Wolf, Ada Tam, Isabel Vanderzee, Xiaokun Wang, James I Andorko, Radhika Narain, Kaitlyn Sadtler, Hongni Fan, Daniela Čiháková, Claude Jourdan Le Saux, Franck Housseau, Drew M Pardoll, Jennifer H. Elisseeff

## Abstract

Synthetic biomaterials and medical devices suffer to varying levels from fibrosis via the foreign body response (FBR). To explore mechanistic connections between the immune response and fibrosis from the FBR, we first analyzed fibrotic capsule surrounding human breast implants and found increased numbers of interleukin (IL)17-producing γδ^+^ T cells and CD4^+^ T_H_17 cells as well as senescent cells. Further analysis in a murine model demonstrated an early innate IL17 response to synthetic implants, mediated by innate lymphoid cells and γδ^+^ T cells, was followed by a chronic adaptive antigen dependent CD4^+^ T_H_17 cell response. Mice deficient in IL17 signaling established that IL17 was required for the fibrotic response to materials and the development of p16^INK4a^ senescent cells. Treatment with a senolytic agent reduced IL17 expression and fibrosis. Discovery of a feed-forward loop between the T_H_17 and senescence response to synthetic materials introduces new targets for therapeutic intervention in the foreign body response.

## Main Text

Synthetic biomaterials serve as the building blocks of medical devices and implants. Biomaterials were historically selected based on their physical properties such as mechanical strength and durability while at the same time inciting minimal host response after implantation. Despite that many advances that medical implants bring to medicine, synthetic materials suffer to varying extents from the foreign body response (FBR) that leads to a capsule of fibrous tissue surrounding the implant [1]. Manipulating chemistry and surface properties can mitigate the FBR to a degree, but even a minor response can lead to device failure over time which necessitates surgical removal. While fibrosis may be leveraged to stabilize some implants such as orthopedic implants or stents, it can also lead to implant contraction in the case of hernia meshes and breast implants. Silicone breast implants are widely used in medical practice but develop fibrotic capsules that can necessitate replacement [2]. Further, some recipients experience breast implant syndrome that includes increased risk of rheumatologic disorders [3]. Recent reports on lymphomas arising around synthetic breast implants designed with a surface to enhance fibrotic immobilization further validate the relevance of murine studies demonstrating the pro-carcinogenic potential of the FBR [4-6].

The classic FBR to synthetic materials was first defined in the 1970s [7-9]. It is characterized by protein adsorption and complement activation followed by migration of pro-inflammatory innate immune system cells, in particular, neutrophils and macrophages. Macrophages fuse to form foreign body giant cells and fibroblasts are activated to secrete extracellular matrix leading to formation of fibrous capsule. Macrophages and the innate immune response are considered central to the FBR and implant fibrosis, however, since the innate and adaptive immune systems are intimately connected, it is possible that the adaptive immune system is also contributing to the FBR [10]. Implantation of a biomaterial or clinical devices may therefore impact immune memory and systemic immune responses with yet unexplored clinical consequences.

T cells are a key component of the adaptive immune system that is increasingly recognized for their role in wound healing and tissue repair. CD4^+^ helper T cells regulate bone, liver, and muscle repair processes [11-13]. The T_H_2 T effector cells responding to pro-regenerative biological scaffolds secrete interleukin 4 (IL4) and direct the function of macrophages to promote muscle repair [13]. The presence of T cells has been recognized in the FBR in animal models and surrounding clinical implants but their nature, activation status and role in the response is still largely unknown [10, 14]. Adaptive responses that depend on T cells, which conventionally recognize MHC-presented peptide antigens, have not been seriously considered in the response to synthetic materials despite their increasing association with fibrotic disease [15-17]. Beyond T cells, the influence of other immune cell types such as gamma-delta (γδ) T cells and innate lymphoid cells (ILCs) in regulation of the biomaterials response, tissue damage, and fibrosis remains unexplored.

The connection between the immune response to biomaterials, fibrosis and senescent cells (SnCs) is also unexplored. SnCs are characterized by growth arrest but are far from quiescent. Accumulation of SnCs is associated with age-related chronic diseases but they also regulate development and wound healing [18, 19]. SnCs exert their effects through the secretion of a senescence associated secretory phenotype (SASP). The SASP includes many immunological cytokines that are being associated with specific immune phenotypes [20]. Clearance of SnCs using knockout models and senolytic drugs reduces fibrosis in idiopathic pulmonary fibrosis (IPF) and reduces inflammation and disease symptoms in arthritis, diabetes, and cardiovascular disease [21-24]. Stromal cells such as fibroblasts, endothelial cells and immune cells have all been found to undergo senescence in various tissues and disease states [25, 26].

Here, we identified adaptive immune regulators of the FBR to synthetic materials. ILCs, γδ^+^ and CD4^+^ T cells were the primary sources of IL17 that promote a fibrotic response to human breast implants and a variety of biomaterials in murine models. We then established the interplay between IL17 and senescent cells as a mechanism linking the chronic immune response to synthetic implants to excessive fibrosis. Therapeutic interventions targeting IL17 and senescence reduce fibrosis and inflammation opening the door to new clinical strategies to address the FBR.

### Interleukin 17 secreted by T cells is associated with fibrosis in tissue surrounding human breast implants

Breast implants suffer from fibrosis that can cause capsular contraction which occasionally necessitates removal and replacement. We performed detailed analysis of the immune cells in multiple tissue samples surrounding implants removed from patients undergoing breast implant exchange surgery. All implants had a silicone shell and were either temporary tissue expanders filled with saline (or air), or permanent implants filled with silicone or saline. Implant samples included both normal and textured surface properties. Peri-implant samples included tissues surrounding implants with both smooth and textured surface properties. Implants were originally placed adjacent to either adipose or muscle tissue depending on pre- or sub-pectoral implantation technique. Average patient age was 56 (range of 41-70 years) and average implant residence time was 41 months (range of 1-360 months). For each implant capsule, we profiled up to 4 tissue sections including left anterior, left posterior, right anterior, and right posterior with respect to the anatomical position of the implant (SFig 1A).

**Figure 1.**
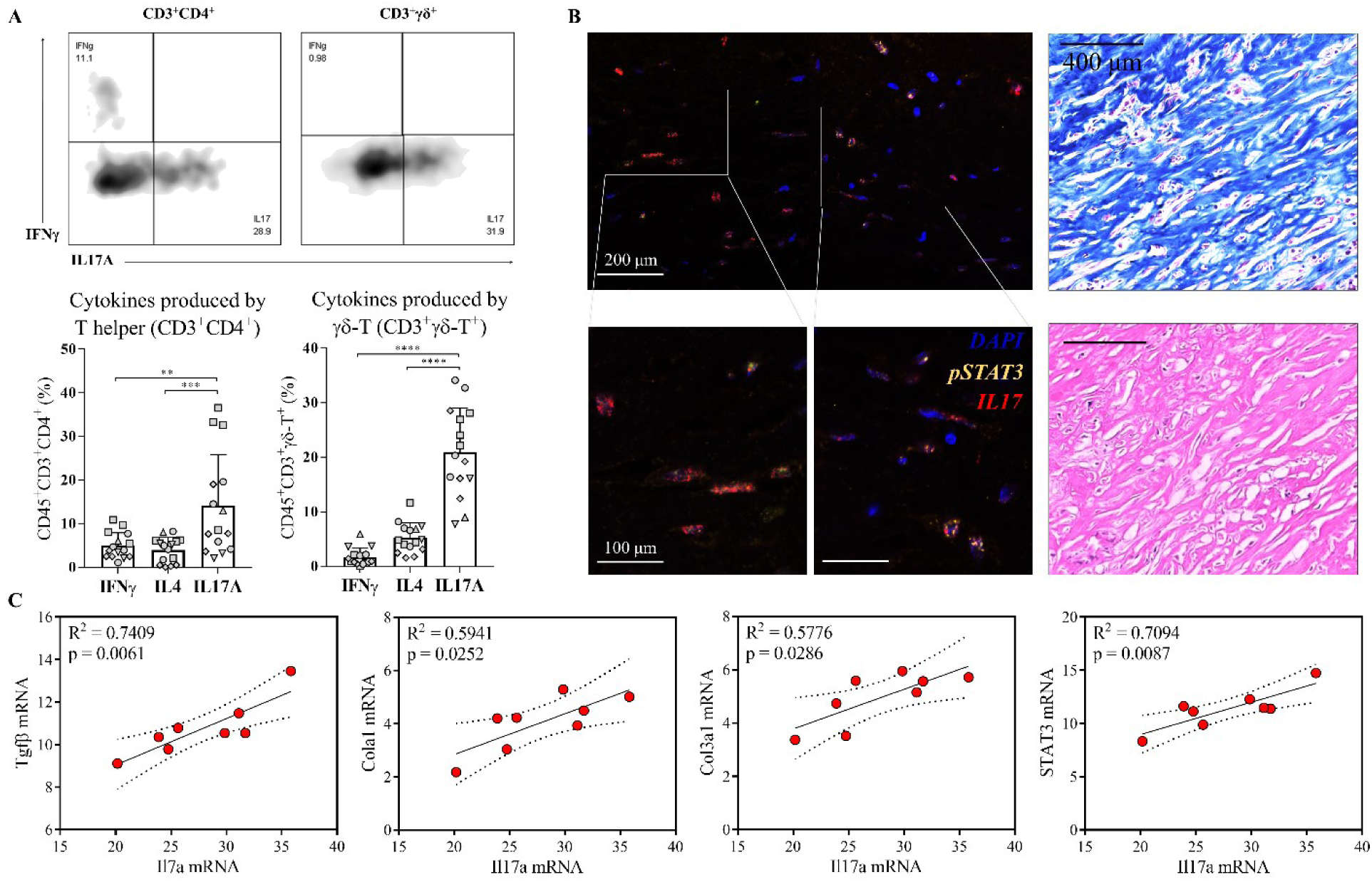
IL17 is secreted by gamma-delta and CD4+ T cells in tissue surrounding human breast implants and correlates with expression of fibrosis markers. Tissue samples surrounding silastic tissue expanders and implants were evaluated by flow cytometry and gene expression analysis. (A) Flow cytometry revealed T helper cells (CD45^+^Thy1.2^+^CD3^+^CD4^+^) T cells and γδ^+^ T cells (CD45^+^Thy1.2^+^CD3^+^γδ^+^) from implant-associated tissue produced IL17A at significantly greater levels compared to interferon gamma (IFNγ), and interleukin 4 (IL4). The percentage of IL17A^+^ cells in both populations was higher compared to IFNγ or IL4 secreting cells. Samples from individual patients are designated with different shapes. (B) Immunofluorescence staining of pSTAT3 (green) and IL17 (red) in the fibrous capsule along Masson’s trichrome and H&E staining. (C) Correlation of qRT-PCR gene expression between Il17a mRNA and fibrosis-associated genes, including collagen I (Col1a1), collagen III (Col3a1), and Tgfβ. Data are means ± SD, n = 3 (A), n = 8 (C) ANOVA [(A) and (C)]: ****P < 0.0001, ***P < 0.001, **P < 0.01, *P < 0.05.

Multiparametric flow cytometry of infiltrating CD45^+^ leukocytes revealed the presence of large numbers of CD3^+^ T cells in addition to myeloid populations of mononuclear phagocytes, dendritic cells, eosinophils, and granulocytes (SFig 1B and C). Intracellular cytokine staining of CD4^+^ T cells revealed significantly higher levels of IL17 producing cells (T_H_17) compared to interferon gamma (IFNγ) (T_H_1) and IL4 (T_H_2)-producing cells in the tissue surrounding the implants (Fig 1A). γδ^+^ T cells (CD45^+^CD3^+^γδ^+^) represented a high proportion of the total CD3^+^ cells (Mean ± SD: 16.97% ± 8.98%, SFig 1E) around the implants and expressed similar levels of IL17 as the CD4^+^ T cells. Immunofluorescence confirmed the presence of IL17 with concomitant nuclear staining of phosphorylated signal transducer and activator of transcription 3 (pSTAT3) that is essential for IL17 expression (Fig 1B, SFig 1D) [27]. Overall, these results support a consistent type 17 immune response to human silicone implants independent of the surface properties and implantation site, with both γδ^+^ and CD4^+^ T cells serving as the primary contributors to IL17 production.

To investigate if the T_H_17/IL17 immune signature contributed to the fibrosis observed in human tissue surrounding the implants (Fig. 1B), we evaluated mRNA levels of collagen I and III, transforming growth factor β (*Tgfβ*), and the fibroblast specific protein *S100a4* (Fig 1C, SFig 1F) [28-30]. Expression of *Il17* positively correlated with expression of *collagen I* (R^2^ = 0.5941, p = 0.0252) and *collagen III* (R^2^ = 0.5776, p = 0.0286). The expression of *Il17* also correlated with the fibrosis-related growth factor *Tgfβ* (R^2^ =0.7409, p =0.0061). There was also a significant correlation between *Il17* expression and *STAT3* (R^2^ =0.7094, p= 0.0087), indicating a STAT3-dependent mechanism, further supporting the relevance of T_H_17 T cells in the FBR in patients. Together, these results suggest that the IL17 produced by γδ^+^ and CD4^+^ T cells may contribute to the regulation of fibrogenesis in response to the breast implants.

### Innate and adaptive lymphocytes sequentially contribute to chronic IL17 production in response to synthetic materials

To further investigate the role of IL17 in implant-associated fibrosis suggested by our clinical findings, and understand the mechanism linking the type 17 immune response to the FBR, we implanted synthetic materials in C57BL/6 mice. Materials were placed both subcutaneous or in a muscle wound to mimic surgical implantation and tissue trauma. Poly(caprolactone), PCL, was used as the primary model synthetic material since it induces a robust inflammatory response and is also a component of clinical implants [31]. Results were validated with additional clinically-relevant synthetic materials including polyethylene (PE), polyethylene glycol (PEG), and silicone.

Implantation of PCL in surgical dermal and muscle wounds increased IL17 expression in the tissue compared to saline controls. We identified three primary cell sources of IL17 that evolved over time: IL17 producing group 3 innate lymphoid cells (ILC3s; CD45^+^CD3^-^Thy1.2^+^) that can respond quickly to danger associated molecular pattern (DAMPs) and cytokines independently of TCR signaling, IL17-producing γδ^+^ T cells (γδ T17) that can recognize non-peptide antigens such as lipids, phosphoantigens and carbohydrates, and adaptive CD4^+^ T cells (T_H_17) that can be activated, specifically through the alpha beta TCR receptor (Fig 2A-D) [32].

**Figure 2.**
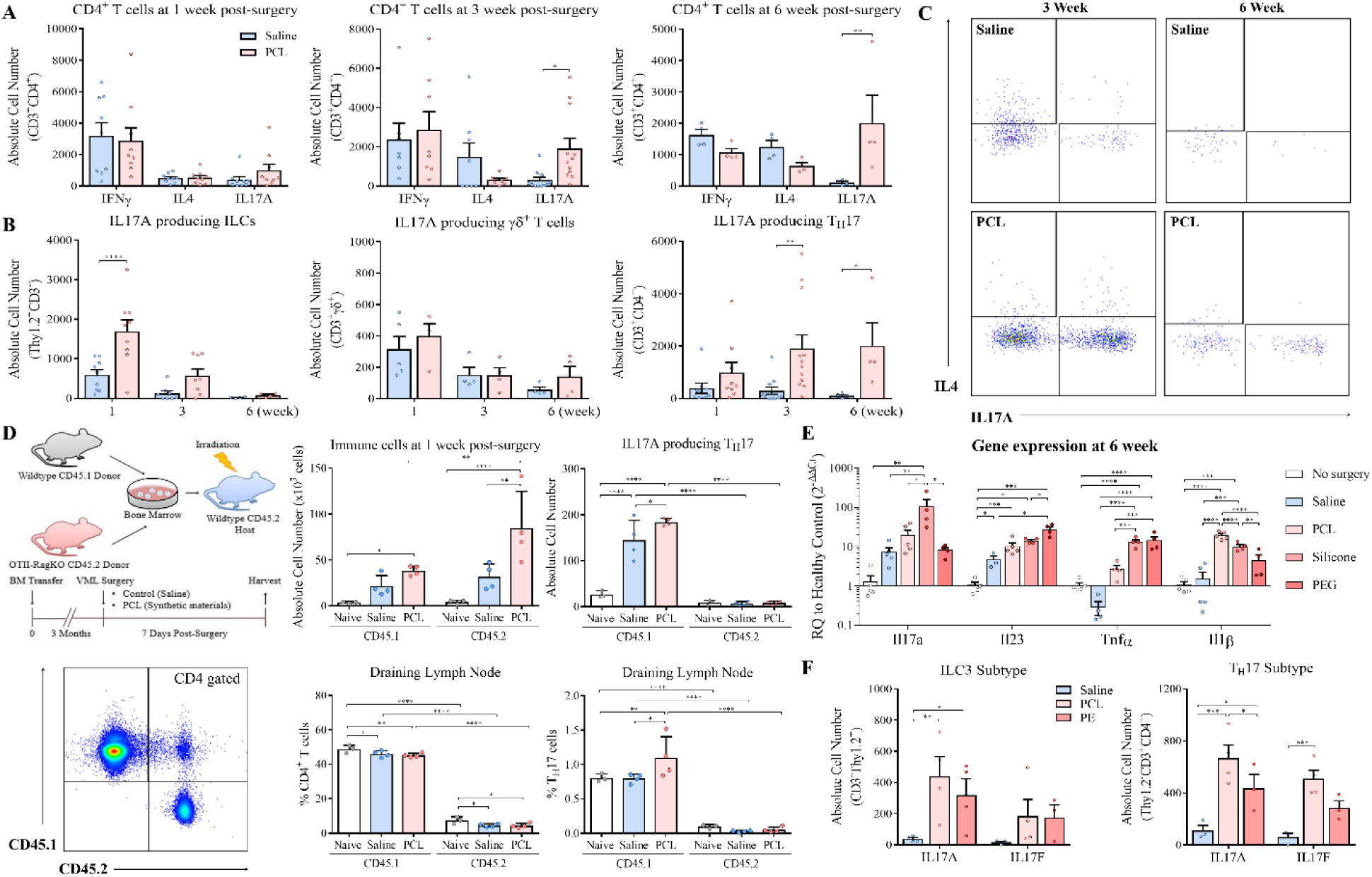
Synthetic materials induce an IL17 immune response. (A) C56BL/6 mice received quadricep muscle injuries and were subsequently implanted with synthetic biomaterial particles or saline. Numbers of IFNγ, IL4, and IL17A secreting T cells were quantified at different time points. While IFNγ and IL4 were unchanged, IL17a significantly increased in mice with synthetic implants compared to saline control treated mice at 3 and 6-weeks post-surgery. (B) Kinetics of IL17A expression by different cell types, including innate lymphoid cells (ILCs), γδ+ T cells, and CD4^+^ T helper cells over time. (C) Representative flow cytometry plots of IL17A production by CD4^+^ T cells at 3 and 6-weeks post-surgery. (D) IL17A and IL17F cytokines were quantified in ILCs and TH17 cells by flow cytometry in muscle implanted with PCL and PE at 3-weeks post-injury. (E) qRT-PCR gene expression of Il17a and other inflammatory genes such as Il1β, Il23, and Tnfα in tissue surrounding PCL, PEG, or silicone implants 6-weeks after implantation. Data are displayed as relative quantification (RQ) to healthy tissue controls. (F) Comparison of IL17A production by ILCs and CD4+ T cells in wildtype and OTII-Rag^-/-^ mice at 1-week post-injury. Data are means ± SD, n = 8 [(A), (B)], n = 5 (D), n = 4 (E). ANOVA [(A), (B), (D), (E), and (F)]: ****P < 0.0001, ***P < 0.001, **P < 0.01, *P < 0.05.

At 1-week post-implantation in the murine model, ILCs and γδ^+^ T cells were the primary source of IL17 (Fig 2B). CD4^+^ T cells exhibited no significant differences between IFNγ, IL4 or IL17 expression at one week by intracellular staining (Fig 2A, 2B). After 3 and 6 weeks, IL17 expression shifted from ILC3s and γδ T17 cells to T_H_17 cells. Both IL17A and IL17F expression significantly increased in T cells (CD3^+^CD11b^-^) sorted from PCL-associated tissue compared to saline control at 1-week post-injury (SFig 3A). Expression of *Rorc* (coding RORγt), *Il23r*, and *Dusp4*, also increased in biomaterial-associated T cells further confirming the type 17 immune profile. Analysis of the immune response to PCL in IL17A-GFP reporter mice validated that CD4^+^ T cells are the primary source of IL17A and there was minimal IL17A expression in myeloid cells at 3-weeks post-surgery (SFig 2D). Finally, gene expression analysis of the whole tissue confirmed significant upregulation of *Il17a* expression in response to PCL implants compared to no implants (saline controls) (SFig 3C). Expression of additional pro-inflammatory cytokines *Il1β*, tumor necrosis factor alpha (*Tnfα*), and *Il23p19* also increased in the tissue (Fig 2E, SFig 3C). These pro-inflammatory cytokines drive immune activation and chronic inflammation through the differentiation and activation of T_H_17 cells [33].

**Figure 3.**
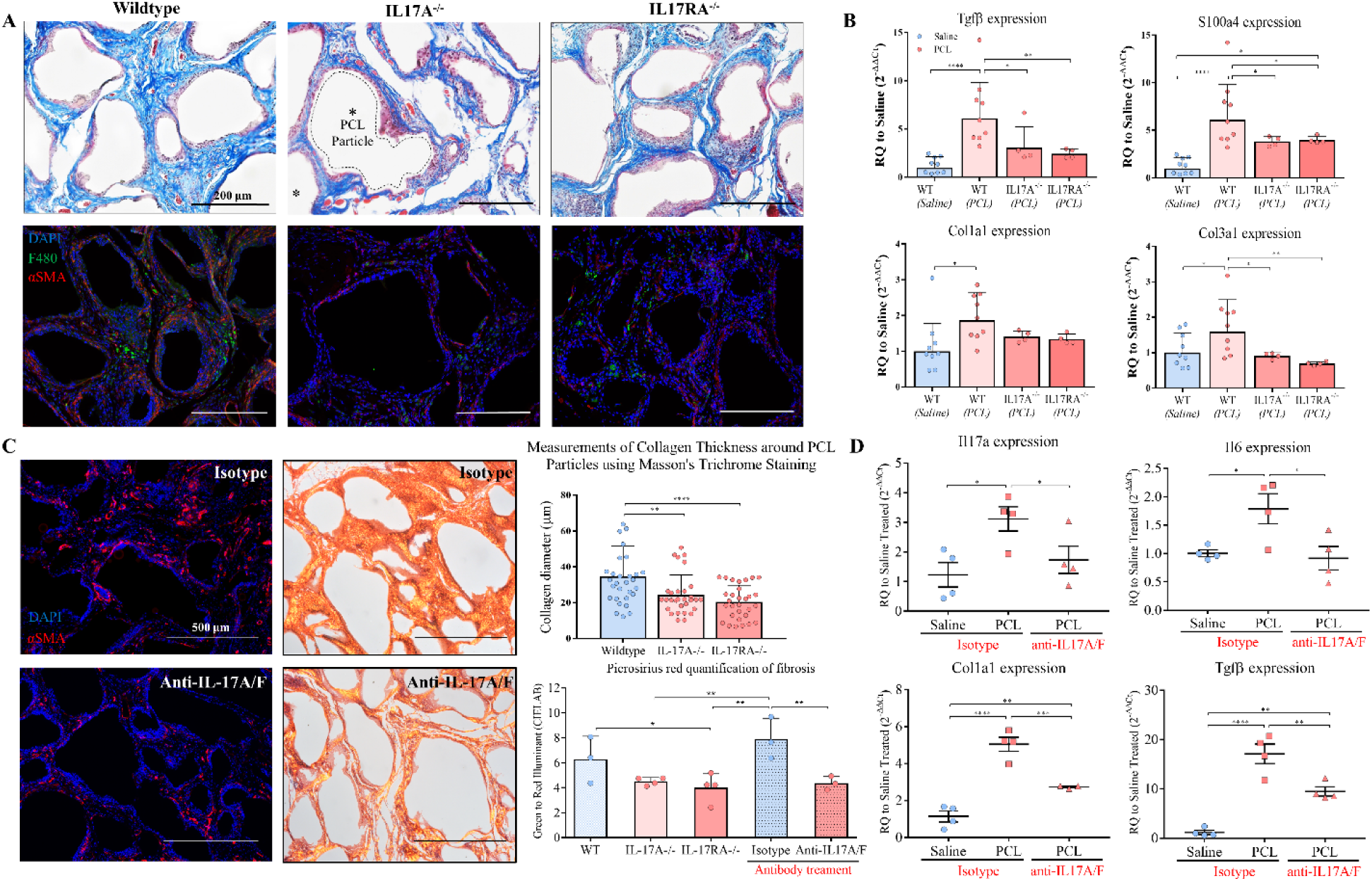
IL17 inhibition reduces the fibrotic response to synthetic materials. PCL was implanted in transgenic mice lacking IL17A expression and IL17 signaling through the IL17RA receptor. (A) Histological staining of implants 12-weeks for collagen (Masson’s trichrome) and immunofluorescence for the macrophage marker F4/80 and fibrosis-associated αSMA protein were evaluated. (B) qRT-PCR gene expressions of fibrotic markers including TGFβ, S100a4, and collagen III were analyzed in WT, IL17A^-/-^, and IL17RA^-/-^ mice. (C) Co-administration of IL17A and IL17F neutralizing antibody (100 μg/mL each) or isotype control (mouse IgG1) was given intraperitoneally to mice with PCL implanted for 4 weeks. To evaluate the degree of fibrosis, tissues was harvested at week 6 for histological assessment. Immunofluorescence staining showed reduced αSMA markers in mice with antibody treatment compared to isotype control. Picrosirius red stain showed the spectrum of color (green to red) relative to the degree of collagen density (thinnest to thickest respectively) and green to red illuminant was quantified by CIELAB. Thickness of the fibrous capsule in WT and IL17KO mice was also measured. (D) qRT-PCR gene analysis of Il17a, Il6, Tgfβ, and type I collagen in tissues treated with neutralizing antibody compared to isotype control at 6-weeks post-injury. Data are displayed as relative quantification (RQ) and saline control. Data are means ± SD, n = 9 (WT mice), n = 4 (IL17 knockout mice), ANOVA [(B) and (C)]: ****P < 0.0001, ***P < 0.001, **P < 0.01, *P < 0.05.

While the degree and nature of the T_H_17 and type 17 immune response varied depending on the biomaterial composition and physical properties, a range of synthetic materials with varying chemistry and physical properties activated this pathway in different tissue environments. Flow cytometry and gene expression analysis confirmed the T_H_17 response to multiple biomaterial types over 6 weeks of implantation. At 3 weeks, flow cytometry showed expression of both IL17A and IL17F in CD4^+^ T cells and ILCs (type 3) in response to PCL and PE, with most cells expressing both forms of the protein (Fig 2F). At 6 weeks, gene expression of type 17 immunity-associated genes *Il1β, Tnfα, Il23*, and *Il17a* increased in response to PCL, silicone, and PEG (Fig 2E). PCL implanted into the subcutaneous space with less tissue disruption or injected directly into muscle without a large wound also activated a type 17 response (SFig 3D). There was elevated IgG1 in the blood of mice implanted with PCL compared to saline controls at 6-weeks post-surgery, highlighting B cell response and further supporting engagement of the adaptive immune system (SFig 4A). These results, combined with the clinical data from breast implants, suggest that the T_H_17 response is activated in response to implants of various size, shape, and texture in different tissue environments.

**Figure 4.**
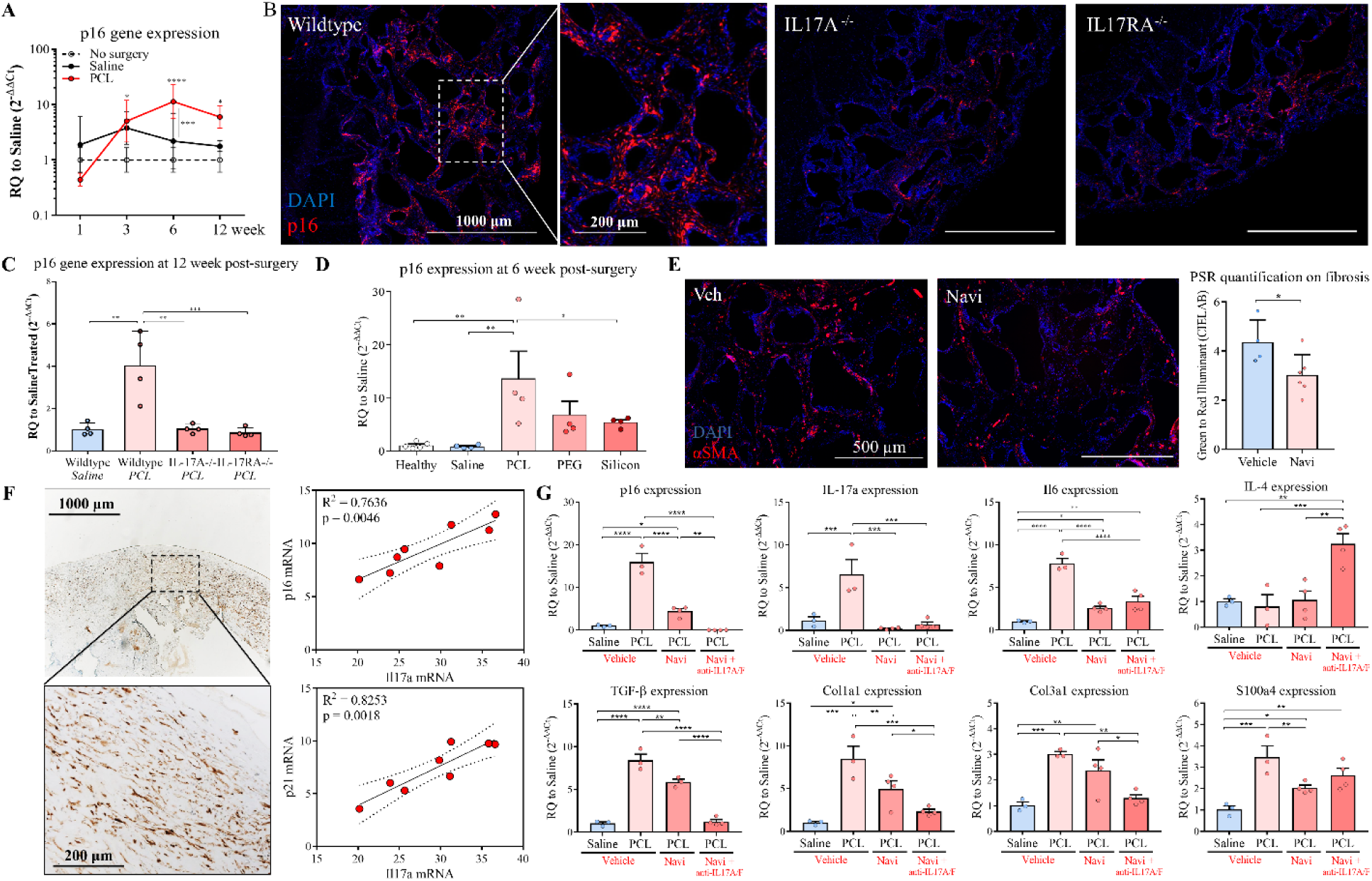
p16^INK4a+^ senescent cells develop around synthetic materials but are abrogated in IL17A^-/-^ and IL17RA^-/-^ mice. (A) qRT-PCR gene expression of p16^INK4a^ progression normalized to healthy tissue control over time. (B) Immunofluorescence staining of p16^INK4a^ (red) in wildtype mice, IL17A^-/-^, and IL17RA^-/-^ mice. (C) qRT-PCR analyses of p16^INK4a^ comparing WT mice to IL17A^-/-^ and IL17RA^-/-^ mice with synthetic implants 12-weeks post-injury. (D) Gene expression of p16^INK4a^ comparing different synthetic materials 6-weeks post-injury. (E) Navitoclax (ABT-263, or Navi) was administrated to eliminate senescent cells. Images showed expression of αSMA comparing to vehicle (DMSO) and Navi treated mice, followed by quantification of G-R illuminant by CIELAB using PSR. (F) Immunohistochemistry images of p16^INK4a^ on human fibrotic capsule. Linear regression showed the correlation between *Il17a* mRNA expression and *p16*^*INK4a*^ and *p21* mRNA. Dotted lines indicated the 95% confidence intervals (CI) of the fitted line. (G) Mice received Navi treatment alone or in combination with IL17A/F neutralizing antibody were harvested at 6-weeks post-injury to evaluate gene expression of *p16*^*INK4a*^, *Il17a, Il6, Il4, Tgfβ, S100a4, collagen I* and *III*. Data are means ± SD, n = 4[(A), (C), (D), (G)], n = 8 (F). ANOVA [(A) (C) (D) (E) and (F)]: ****P < 0.0001, ***P < 0.001, **P < 0.01, *P < 0.05.

To determine whether the T_H_17 response was mediated by the TCR, we used wildtype CD45.1 and OTII-Rag^-/-^ CD45.2 bone marrow chimera mice (Fig 2D). The OTII TCR transgenic Rag^-/-^ CD4^+^ T cells are specific of ovalbumin (OVA) so we can test if there is non-specific activation of OVA-specific T cells in the context of a highly diverse TCR repertoire of the CD45.1 cells in the wound environment with or without PCL implantation. We found that CD45.1^+^ and CD45.2^+^ immune cells from donors responded to PCL implantation after 1 week. In the tissue and draining lymph nodes CD4+ T cells were primarily from the WT CD45.1 donor (Fig 2D). IL17 was expressed in wildtype CD45.1 CD4^+^ T cells in response to PCL implantation but was absent in the CD45.2 OTII-Rag^-/-^ T cells. The capacity of OTII-Rag^-/-^ T cells to undergo T_H_17 differentiation was confirmed *in vitro*, so the failure to produce IL17 in response to synthetic implants did not represent an intrinsic defect in their ability to undergo differentiation to T_H_17 (SFig 4B). At 3-weeks post-surgery, CD8^+^ T cells, predominantly produced IFNγ^+^, resolved whereas CD4^+^ T cells remained and continued to produced IL17 (Fig 2B, SFig 2E). This suggests that the IL17 production by CD4^+^ T cells in response to PCL implantation was antigen specific. As previously described, biomaterials can modulate the T cell response to antigens thereby acting as an adjuvant. Spleenocytes from OTII mice cultured *in vitro* with OVA in combination with PCL or PE for 48 hours showed increased T cell proliferation compared to without biomaterials.

### Chronic IL17 in response to synthetic implants promotes fibrosis

T_H_17 immune responses are implicated in fibrotic diseases in multiple tissue types including skin, heart, lung, and liver, though have not been studied in the context of FBR [34-37]. In the wild type (WT) mice that demonstrated a type 17 immune response, the expression of fibrosis related genes after surgery evolved over time (SFig 4D). Without a biomaterial, expression of the extracellular matrix genes collagen I and III initially increased after surgery then decreased by 6 weeks as normal healing progressed and tissue repaired. The presence of the PCL implant significantly increased expression of collagen III and the fibrosis gene S100a4 after 6 and 12 weeks. Masson’s trichrome and immunohistochemistry confirmed increased collagenous extracellular matrix and alpha smooth muscle actin protein (αSMA) surrounding the PCL particles (Fig 3A), features typical of the FBR and fibrotic capsule formation.

To evaluate if fibrosis associated with the FBR to biomaterials was IL17 dependent, we compared the implant response in *Il17a*^-/-^ and *Il17ra*^-/-^ mice with WT mice. We found that expression of fibrosis-related genes *S100a4, Tgfβ, collagen I* and *III* significantly decreased in both knockout mice implanted with PCL compared to the WT mice (Fig 3B). The quantity, composition, and organization of ECM visible around the PCL particles decreased in the knockout animals (Fig 3A). αSMA immunofluorescence was absent in *Il17* signaling deficient and picrosirius red (PSR) staining significantly shifted. PSR visualized with a polarized lens produces birefringence that is specific for collagen fiber organization; larger collagen fibers are bright orange to red and thinner fibrils are green to yellow [38]. A reduction in collagenous matrix and thinner fibrotic capsules around the PCL particles was confirmed in knockout mice compared to WT mice at 12-weeks post-surgery using image analysis and quantification of collagen thickness and green to red illuminant on PSR images (Fig 3C). In addition, expression of the inflammasome associated gene Nlrp3 and Il1β significantly decreased in PCL-implanted tissue in IL17raKO mice, but not in IL17aKO, compared to the WT mice (SFig 5B). These findings demonstrate that impaired IL17 signaling may reduce fibrosis that occurs in response to PLC implantation.

We next assessed the myeloid response associated with PCL implantation in the IL17 knockout mice. The number of neutrophils and macrophages significantly increased in tissue implanted with synthetic materials. Macrophages sorted from the tissue implanted with PCL expressed higher levels of profibrotic stimulating factors that directly activate fibroblasts including *Tgfβ*, platelet derived growth factor α (*Pdgfα*), and vascular endothelial growth factor α (*Vegfα*) (SFig 6B). While the number of monocytes in the IL17A^-/-^ and IL17RA^-/-^ mice did not change, the proportion of MHCII^high^ and MHCII^low^ macrophages was significantly altered. MHCII^high^ macrophage numbers decreased to levels of the WT controls without implants in the IL17RA^-/-^ mice while MHCII^low^ macrophages significantly decreased in both IL17A^-/-^ and IL17RA^-/-^ mice. F4/80 immunofluorescence confirmed decreased macrophage recruitment to the implants in the IL17 knockout animals (SFig 6D). Neutrophils (CD11b^+^Ly6c^+^Ly6g^+^) and macrophages (CD11b^+^F4/80^+^), decreased in the IL17A^-/-^ and IL17RA^-/-^ mice compared to WT after 12 weeks (SFig 6C, 6D). Flow cytometry and immunofluorescence confirmed fewer neutrophils in the IL17A^-/-^ and IL17RA^-/-^ mice implanted with PCL. Functional recovery of the tissue repair was not impaired in the IL17A^-/-^ and IL17RA^-/-^ mice as assessed by treadmill testing (SFig 4C).

Results from the knockout studies suggest that blocking of IL17 signaling may reduce FBR-associated fibrosis without negatively impacting healing. We therefor sought to determine the therapeutic potential of blocking IL17 on fibrosis. Since IL17A and IL17F were both produced in response to the implant and the IL17RA^-/-^ mice showed the greatest reduction in fibrosis compared to IL17A^-/-^, we evaluated delivery of IL17A and IL17F specific neutralizing antibodies (IL17A/F NAbs) to test the impact of functional neutralization of IL17R signaling on fibrosis. Mice were treated with IL17A/F NAbs 4 weeks after biomaterial implantation every other day for one week for a total of 5 injections. One week after the IL17A/F NAb treatment, *IL17* and *IL6* expression decreased. Expression of fibrosis-associated genes *Tgfβ* and *Collagen I* also decreased (Fig 3D). Immunofluorescence of αSMA also decreased, further suggesting reduced fibrosis. Histologically, collagen density and fiber organization decreased with anti-IL17A/F treatment compared to isotype controls (Fig 3C). PSR staining and quantification demonstrated that IL17A/F NAbs modulated ECM organization and reduced fibrosis to levels that were similar to the knockout animals.

### p16^INK4a^ senescent cells develop during the FBR and senolysis reduces fibrosis

IL6 is a critical mediator contributing to the differentiation of T_H_17 cells. Since IL6 is associated with the SASP produced by senescent cells, we investigated whether SnCs were a potential source of IL6 contributing to the chronic IL17 production and fibrosis [18]. SnCs are characterized by expression of *p16*^*INK4a*^, *p21* and SASP factors [39-41]. Consistent with the kinetics of fibrosis development, *p16*^*INK4a*^ expression increased significantly from 6 to 12 weeks after PCL implantation compared to saline controls (Fig 4A). *p16*^*INK4a*^ positive cells were localized to the peri-implant region and exhibited a fibroblastic morphology with a single nucleus (Fig 4B, SFig 7A). Fibroblasts sorted from PCL implants expressed significantly higher levels of *p16*^*INK4a*^ compared with healthy and saline controls, whereas sorted macrophages did not express *p16*^*INK4a*^ (SFig 6B, 7B). All materials tested induced *p16*^*INK4a*^ expression to varying levels, again suggesting that the response is material independent (Fig 4D). SnC development was abrogated in both the IL17A^-/-^ and IL17RA^-/-^ mice at 12 weeks to levels of the no implant controls, demonstrating that their development depended on IL17 (Fig 4C).

To further validate and define clinical relevance for the association of SnCs with fibrosis and the FBR, we evaluated patient tissue from breast implant exchanges. Significant numbers of *p16*^*INK4a*^ positive cells were present in the tissue surrounding the implants (Fig 4F). Gene expression analysis of tissue surrounding the implants also showed a strong positive correlation between *Il17* and *p16*^*INK4a*^ (R^2^ = 0.7636, p = 0.0046) in addition to *Il17* and *p21* (R^2^ = 0.8253, p = 0.0018, Fig 4F).

To determine if clearance of SnCs associated with the FBR could reduce fibrosis, we administrated the senolytic agent Navitoclax (Navi, ABT-263) that selectively kills SnCs [25, 42]. Navi was administered alone and in combination with anti-IL17A/F NAbs 4 weeks after implantation, when *p16*^*INK4a*^ expression significantly increased with biomaterial implantation compared to control (Fig 4A). One week after treatment (6 weeks after implantation), expression of *p16*^*INK4a*^ significantly decreased confirming senolysis (Fig 4G). Clearance of the SnCs reduced αSMA immunofluorescence, suggesting reduced fibrosis activity (Fig 4E). Concomitant with p16^INK4a^ and αSMA reduction, *Il17a, Il6, Tgfβ, S100a4, collagen I* and *III* gene expression decreased after senolytic treatment. These genes decreased even further when the senolytic was co-administered with anti-IL17A/F NAbs (Fig 4G). Altogether these findings demonstrate that cellular senescence sustains excess fibrosis and links to chronic IL17 and fibrogenesis during the foreign body response.

## Discussion

Medical implants derived from synthetic materials suffer to some extent from the FBR that not only impairs implant function and longevity but may also in turn impact the host. There are numerous anecdotal reports of systemic illnesses associated with orthopedic and soft tissue implants most notably the breast implant syndrome [43, 44]. With only local inflammatory components considered relevant to the FBR, direct correlation of implant responses with systemic immune pathologies was difficult to connect. Preclinical and clinical observations noted the presence of T cells with implants but their relevance to the fibrotic inflammatory response was unclear [14, 45]. Here we present evidence of an IL17 inflammatory response and cellular senescence in the FBR to biomaterials and implicate them in the associated fibrotic response.

T cells are increasingly recognized for their role in determining repair pathways after tissue damage. We observed a T_H_17 response to biomaterials with multiple implant chemistries in numerous anatomical locations. B cells and their antibody production have already been associated with tissue damage and more recently with synthetic materials [46-48]. For example, a B cell response to synthetic materials was discovered with implantation in the intraperitoneal cavity. Moreover, the response was independent of material chemistry and physical properties [48]. In our studies, we utilized particles of materials in order to fill tissue defects in the murine model. These particulates had different sizes, textures and surface properties and can themselves activate a rapid, inflammasome-mediated innate immune responses [49]. The presence of T_H_17 cells in all of clinical samples tested, from large (non-particulate) and diverse implant surfaces, suggests the IL17 pathway is conserved in biomaterial-associated fibrosis and is broadly relevant and clinically applicable. Further studies may elucidate unique aspects of the IL17 and immune response with different materials.

The induction of type 17 inflammation has been implicated in tissue fibrosis. For example, IL17 is a central contributor to pathogenic lung and liver fibrosis [50, 51]. While T_H_17 responses are a mechanism to combat extracellular pathogens, they are also associated with autoimmune diseases. For example, T_H_17 cells are implicated in rheumatoid arthritis and Sjogren’s syndrome and therapies inhibiting IL17 ameliorate disease symptoms. Induction of a T_H_17 immune response to biomaterials is supported by the chronic neutrophil response to synthetic materials since IL17 induces chronic neutrophilia. Multiple studies have now demonstrated a negligible impact of neutrophil depletion on fibrosis around biomaterials, suggesting that they are not inducing fibrosis but a byproduct [48, 52].

The discovery of senescent cells in the FBR presents a novel mechanism for the sustained type 17 inflammation and fibrosis around implants in addition to a new therapeutic target. Senolytic compounds are being developed to treat numerous age-related diseases including arthritis, cardiovascular and Alzheimer’s disease [53, 54]. Clinical studies testing senolytics in idiopathic pulmonary fibrosis are already showing efficacy and more clinical studies are ongoing [55, 56]. The SASP secreted by SnCs include cytokines associated with a T_H_17 immune response including IL6, IL1β, and the loss of SnCs in the IL17 knockout models further supports the IL17-SnC connection. The morphology of the p16^INK4a^ positive cells appeared fibroblastic and sorted fibroblasts expressed significantly higher levels of p16^INK4a^ suggesting that stromal fibroblasts are one source of senescence in the FBR. It is possible the T_H_17 response is being driven locally by material auto-antigens or antigens, possibly facilitated by the senescent cells, as appears to be the case in post-traumatic osteoarthritis [57].

Canonical activation of T cells requires presentation of antigen and costimulatory factors. The lack of CD4^+^ T cell response to PCL in the chimeric OTII-Rag^-/-^ T cells confirms that the T_H_17 implant response is antigen dependent. One explanation of an antigen-specific T cell response to materials that do not contain protein is a low-level chemical derivatization self-protein by components of the synthetic material, such that they now appear as “non-self”. The finding of IgG in response to implantation of a synthetic material supports this hypothesis, since haptenization of proteins is necessary for T cell help for Ig switching in activated hapten-specific B cells [58, 59]. Combinations of a biomaterial with host proteins can activate T cells as in the case of titanium implant debris that creates a metal-protein complex that can elicit cell responses [60]. Addition of an implant in the context of tissue damage may modulate the natural adaptive damage response by modifying self-antigens or impacting antigen presentation. Finally, there is evidence that T cells can specifically respond to non-peptidic repeating structures such as sugars and lipids and thus may recognize repeating polymer structures [61, 62]. Regardless of the antigen or combination of antigens, the type 17 immune response is activated and directing key aspects of the FBR and targeting this pathway has therapeutic benefits for biocompatibility.

Engagement of the adaptive immune system introduces the potential influence of environment factors in the FBR beyond the standard genetic that can modulate adaptive immune responses such as infection, history of antigen exposure, and the microbiome. Future studies will investigate the impact of these immunomodulatory environmental factors on the T_H_17 and senescence responses to biomaterial implants.

## Acknowledgements

The authors gratefully acknowledge S. Ganguli for assistance with NanoString. We thank YR. Gonzalez and KR. Wagner for help with the treadmilling testing. This work was funded in part by the Bloomberg∼Kimmel Institute, Morton Goldberg Professorship and the Department of Defense.

## Author contributions

L.C, J.H.E, D.P, and F.H conceptualized this study and wrote the manuscript with input from all co-authors. L.C, J.H.E, D.M, A.L contributed to experimental design and interpretation of results. L.C, D.M, A.M, I.V, R.N, M.T.W, X.Wang, J.I.A contributed to conducting experimental procedures. G.R provided human surgical samples. L.C, X.Wu, and H.F assisted with animal breeding and genotyping. C.J.L.S, D.C, and K.S discussed the results and commented on the manuscript.

## Conflict of Interests

COI is provided in attached documentation.

