## Supplemental Data for "Interleukin-17 and senescence regulate the foreign body response"

**Materials and Methods:**

***Clinical samples:*** Tissue was acquired from patients undergoing implant exchange or replacement surgeries (n = 12, JHU IRB exemption IRB00088842). For each patient, up to 4 tissue sections were profiled including the left anterior, left posterior, right anterior, and right posterior with respect to the anatomical position of the implant (SFig 1A). Each section was weighed and 0.25 g of tissue was dissected for histology. Remaining tissue was analyzed depending on the available quantity; qRT-PCR assay (<1 g), flow cytometry (1-2 g), or both (>2 g). In this study, 8 samples were utilized for qRT-PCR and 5 samples were used for flow cytometry from different individuals (n=12). Peri-implant samples included tissues surrounding implants with both smooth and textured surface properties. All implants had a silicone shell and were either temporary tissue expanders filled with saline or air or permanent implants filled with silicone or saline. Average patient age was 56 years old (range of 41-70 years old) and the average implant residence time was 41 months (range of 1-360 months).

***Surgical procedures and implantation*:** All animal procedures were performed in accordance with an approved JHU IACUC protocol. Female mice were aged six to eight weeks-old. Several strains were utilized, including wild type C57BL/6j (Jackson Laboratories), Rag2/OT-II (Taconic), IL17A-/- and IL17RA-/- (courtesy of Dr. Yoichiro Iwakura, University of Tokyo, Tokyo, Japan and Dr. Tomas Mustelin, Amgen, Seattle, respectively). Defects in muscle for material implantation were created as previously described [1]. The resulting bilateral muscle defects were filled with 30 mg of a synthetic material.

Synthetic materials tested include polycaprolactone (PCL) (particulate, $M_n$ = 50000 g/mol, mean particle size < 600 μm, Polysciences), ultrahigh molecular weight polyethylene (PE) (particulate, $M_n$ = 5000k g/mol, mean particle size = 150 μm, Goodfellow), medical graded Silicone (sheet, Summit Medical), polyethylene glycol (PEG) hydrogels. PEG hydrogels were made with polyethylene glycol-diacrylate (PEG-DA, $M_n$ = 3400). PEG-DA (Polysciences) in PBS at 10% w/v concentration was mixed with photoinitiator (Irgacure 2959 solubilized to 10% w/v in 70% ethanol) to a final concentration of 0.05%. A 50 μL volume solution was cast onto each well of a flat bottom 96 wells plate. The solution was photocrosslinked via UV light for 30 mins to form a hydrogel sheet. Both silicone and PEG hydrogels were morselized into particles with a scalpel and placed into the defect. Control implants were injected with 50 μL of PBS (as a no implant control). All materials were UV sterilized prior to use. Directly after surgery, mice were given subcutaneous carprofen (Rimadyl®, Zoetis) at 5 mg/kg for pain relief. Mice were euthanized, and their implants were extracted at different time points: 1, 3, 6, or 12-weeks post-surgery. Additional modes of implantation were subcutaneous and intra-muscular injection. Briefly, synthetic materials were implanted subcutaneously (30 mg) on both flanks of the mice, or injected directly into the quadricep muscles through a 16 G syringe (5 mg of materials).

***Bone marrow chimera*:** Three million bone marrow cells from each donor mouse (CD45.1 WT mice and CD45.2 OTII-RagKO mice) were injected intravenously into lethally irradiated (1200 cGy: twice at 600 cGy at 4 hours intervals) to 6-weeks old recipients. Bone marrow reconstituted mice were rested for 3 months prior to the VML procedure.

***Gene expression analysis*:** Total RNA was isolated from whole tissue using TRIzol reagent and Qiagen's RNeasy kits. qRT-PCR was performed using Power SYBR Green Master Mix (Applied

Biosystems) or TaqMan Gene Expression Master Mix (Applied Biosystems) according to manufacturer's instructions. Briefly, 2 µg of total RNA was used to synthesize cDNA using Superscript IV VILO Master Mix (ThermoFisher Scientific). The cDNA concentration was set to 45 ng/well (in a total volume of 20 µL PCR reaction). The qRT-PCR reactions were performed on the StepOne Plus Real-Time PCR System (ThermoFisher Scientific) using manufacturer recommended settings for quantitative and relative expression. All analyses of qRT-PCR data utilized the Livak Method, wherein ΔΔCt values are calculated, and then reported as relative quantification values calculated by $2^{-\Delta\Delta Ct}$ [2]. For tissue samples, *β2m* was used as the reference gene, and samples were normalized to saline treated controls.

SYBR Green Mouse mRNA Primers:

| Primer | Forward Sequence | Primer | Reverse Sequence |
|---|---|---|---|
| β2m forward | CTC GGT GAC CCT GGT CTT TC | β2m reverse | GGATTT CAA TGT GAG GCG GG |
| Tnfα forward | GTC CAT TCC TGA GTT CTG | Tnfα reverse | GAA AGG TCT GAA GGT AGG |
| Il1β forward | GTA TGG GCT GGA CTG TTT C | Il1β reverse | GCT GTC TGC TCA TTC ACG |
| Retnla forward | CTT TCC TGA GAT TCT GCC CCA G | Retnla reverse | CAC AAG CAC ACC CAG TAG CA |
| Ifnγ forward | TCA AGT GGC ATA GAT GTG GAA | Ifnγ reverse | TGA GGT AGA AAG AGA TAA TCT GG |
| Il4 forward | ACA GGA GAA GGG ACG CCA T | Il4 reverse | ACC TTG GAA GCC CTA CAG A |
| Il17a forward | TCA GCG TGT CCA AAC ACT GAG | Il17a reverse | CGC CAA GGG AGT TAA AGA CTT |
| Arginase 1 forward | CAG AAG AAT GGA AGA GTC AG | Arginase 1 reverse | CAG ATA TGC AGG GAG TCA CC |
| Cdkn2a (p16) forward | AAT CTC CGC GAG GAA AGC | p16 reverse | GTC TGC AGC GGA CTC CAT S |
| Cdkn2b (p15) forward | AGA TCC CAA CGC CCT GAA C | p15 reverse | CCC ATC ATC ATG ACC TGG ATT |
| l10 forward | CAG GAC TTT AAG GGT TAC TTG GGT | Il10 reverse | GCC TGG GGC ATC ACT TCT AC |

Mouse TaqMan gene expression primers:

| Primer | Assay ID: | Primer | Assay ID: |
|---|---|---|---|
| β2m | Mm00437762 | Il17a | Mm00439618 |

| | | | |
|---|---|---|---|
| Col1a1 | Mm00801666 | Il23a | Mm00518984 |
| Col3a1 | Mm00802300 | S100a4 | Mm00803372 |
| Pai1 | Mm00435858 | Snai1 | Mm00441533 |
| Tgfβ-1 | Mm01178820 | p16 | Mm00494449 |

Human TaqMan gene expression primers:

| Primer | Assay ID: | Primer | Assay ID: |
|---|---|---|---|
| β2m | Hs00187842 | Il17a | Hs00174383 |
| Col1a1 | Hs00164004 | STAT3 | Hs00374280 |
| Col3a1 | Hs00943809 | S100a4 | Hs00243202 |
| p16 | Hs00923894 | Tgfβ-1 | Hs00998133 |
| Cdkn1a (p21) | Hs00355782 | Ifnγ | Hs00989291 |
| Il4 | Hs00174122 | Il6 | Hs00174131 |

***Histopathology:*** Tissues were harvested and fixed in 10% neutral buffered formalin for 24 hours before step-wise dehydration in EtOH, cleared with xylenes, and embedded in paraffin. Samples were sectioned as 7 µm slices using a Leica RM2255 microtome. Samples were stained for histopathological examination using Masson's trichrome (Sigma-Aldrich), hematoxylin and eosin (Sigma-Aldrich), and picrosirius red (Abcam) stains according to standard manufacturer protocols.

Immunofluorescence staining for IL17 and p16 was conducted using tyramide signal amplification (TSA) method with Opal-570 and Opal-650 respectively. Briefly, after blocking with bovine serum albumin, the first primary antibody was incubated at room temperature for 30 mins, followed by 10 mins of incubation with HRP polymer conjugated secondary antibody, and 10 mins of Opal-650. Unbound antibodies were stripped by microwaving in citrate buffer for 15 mins to allow introduction of the next primary antibody (with different Opal dyes). Slides were then counterstained with DAPI for 5 mins before being mounted using DAKO mounting medium. Imaging of the histological samples was performed on a Zeiss Axio Imager A2 and Zeiss AxioVision software ver. 4.2. Subsequent images were stitched together using ImageJ2 software.

***Flow Cytometry:*** Tissue samples were obtained by cutting the quadriceps femoris muscle from the hip to the knee. Tissues were finely diced and digested for 45 min at 37ºC with 1.67 Wünsch U/ml Liberase TL (Roche Diagnostics) and 0.2 mg/ml DNase I (Roche Diagnostics) in RPMI 1640 medium (Gibco). The digested tissues were ground through 100 µm cell strainers (ThermoFisher Scientific) with excess RPMI, and then washed twice with 1X DPBS. Percoll (GE Healthcare) density gradient centrifugation was used to enrich the leukocyte fraction and remove debris from the muscle samples. The enriched cells were washed, and stained with the following antibody panels:

Myeloid panel:

| Fluorophore | Marker | Manufacturer |
|---|---|---|
| Fixable Yellow | Live/Dead | Life Technologies |
| AF488 | MHC-II | BioLegend |
| PerCP-Cy5.5 | CD11b | BioLegend |
| APC-Cy7 | CD11c | eBioscience |
| PE-594 | Siglec F | BioLegend |
| Pacific Blue | Ly6c | BioLegend |
| AF647 | Ly6g | BioLegend |
| V500 | CD45 | BioLegend |
| PE-Cy7 | F4/80 | BioLegend |

For intracellular staining, cells were stimulated for 4 hours with Cell Stimulation Cocktail (plus protein transport inhibitors) (eBioscience) diluted in complete culture media (IMDM supplemented with 5% fetal bovine serum). Cells were washed and surface stained, followed by fixation/permeabilization (Cytofix/Cytoperm, BD), and then stained for intracellular markers. Flow cytometry was performed using Attune NxT Flow Cytometer (ThermoFisher Scientific).

Lymphoid panel:

| Fluorophore | Marker | Manufacturer |
|---|---|---|
| Fixable Near-IR | Live/Dead | Life Technologies |
| BV786 | CD3 | BioLegend |
| FITC | CD4 | BioLegend |
| BV711 | CD8 | BioLegend |
| PE-Texas Red | γδ-TCR | BioLegend |
| APC | IFNγ | BioLegend |
| PE | IL4 | BioLegend |
| AF700 | IL17A | BioLegend |
| V500 | CD45 | BioLegend |
| PE-Cy7 | Thy1.2 | BioLegend |

***Treadmill Testing:*** Prior to testing, mice were trained on a treadmill apparatus at 5 m/min for 5 mins. Mice were run to exhaustion starting at the speed of 5 m/min, with 1 m/min speed increase every minute. Exhaustion was determined when the mouse stopped running, and stayed on the pulsed shock grid for a continuous 30 seconds. All mice were evaluated at 3, 6, and 12-weeks post-injury.

***IL-17 neutralization and senolytic treatment:*** Mice received 100 µl intraperitoneal (IP) injections of anti-IL-17A (100 µg/ml) and anti-IL-17F (100 µg/ml) (provided by Amgen) or isotype control (rat IgG2a) every other day for a week. For senolytic treatment, mice received IP and intramuscular (IM) injections of Navitoclax (100 mg/kg, Selleckchem) for 5 consecutive days. Co-administration of anti-IL17A/F and senolytic treatment were also evaluated. All mice received treatments at 4-weeks post-surgery for a total of 5 injections (1 injection per day per mouse), and were harvested in the following 2 weeks.

***Statistical Analysis:*** The qRT-PCR data are displayed as the mean ± standard deviation (SD) of the RQ values as determined by the Livak Method, calculated by $2^{-\Delta\Delta Ct}$. Treatment data points are normalized to either healthy or saline treated controls. β2M was utilized as the reference gene. In the flow cytometry reports, the total number of cytokine-secreting cells were computed using FlowJo software, and are displayed as the means ± SEM. Two-way ANOVAs were performed using GraphPad Prism v6, with statistical significance designated at $p \leq 0.05$.

***Antibody Titer:*** PCL particles were dissolved in chloroform to coat the bottom of 96-well plates (20 mg/ml). Sera collected 3, 6, and 12-weeks post-injury were serially diluted (in the range of 1:50 to 1:102400) in ELISA Assay Diluent (BioLegend). Prior to loading, the PCL-coated plate was blocked with the assay diluent for 1 hour. After blocking, each dilution of the serum sample was loaded into the plate and incubated for 2 hours. After washing, biotin anti-mouse IgG1, or IgM, or IgA (BioLegend) were added to capture the bound antibody for 1 hour, followed by washing. Streptavidin solution (BioLegend) was added to the wells and incubated for 30 mins. TMB Substrate (BioLgend) was used for HRP detection, and stopped with $H_2SO_4$. Absorbance was read at 450 nm and 570 nm (background control).

***$T_H$17 in-vitro skewing assay:*** Naïve $CD4^+$ T cells were isolated from mouse spleens and lymph nodes using the Miltenyi Biotec Naïve $CD4^+$ Isolation Kit according to manufacturer's protocol. Cells were differentiated using CellXVivo Mouse $T_H$17 Cell Differentiation Kit (R&D Systems) according to manufacturer's protocol. Cells were cultured for 5 days (refreshed the differentiation media at day 3). On day 5, cells were stimulated with Cell Stimulation Cocktail for 4 hours prior to cytokine staining, and examined using flow cytometry.

***Cell proliferation assays***: primary spleenocytes were isolated from OT-II transgenic mice. Cells were labeled with Cell Trace Violet Cell Proliferation Kit (Invitrogen) according to manufacturer's protocol. Three million cells/well were seeded into 6-well plates and were grown in RPMI supplemented with 10% fetal calf serum (Gibco), nonessential amino acids, sodium pyruvate (Invitrogen), and penicillin-streptomycin (Invitrogen). At the same time, OVA and biomaterials (PCL or PE – 10 mg/well) were introduced into culture. After 48 hours, cells were harvested and analyzed using flow cytometry.

### Supplemental Figures

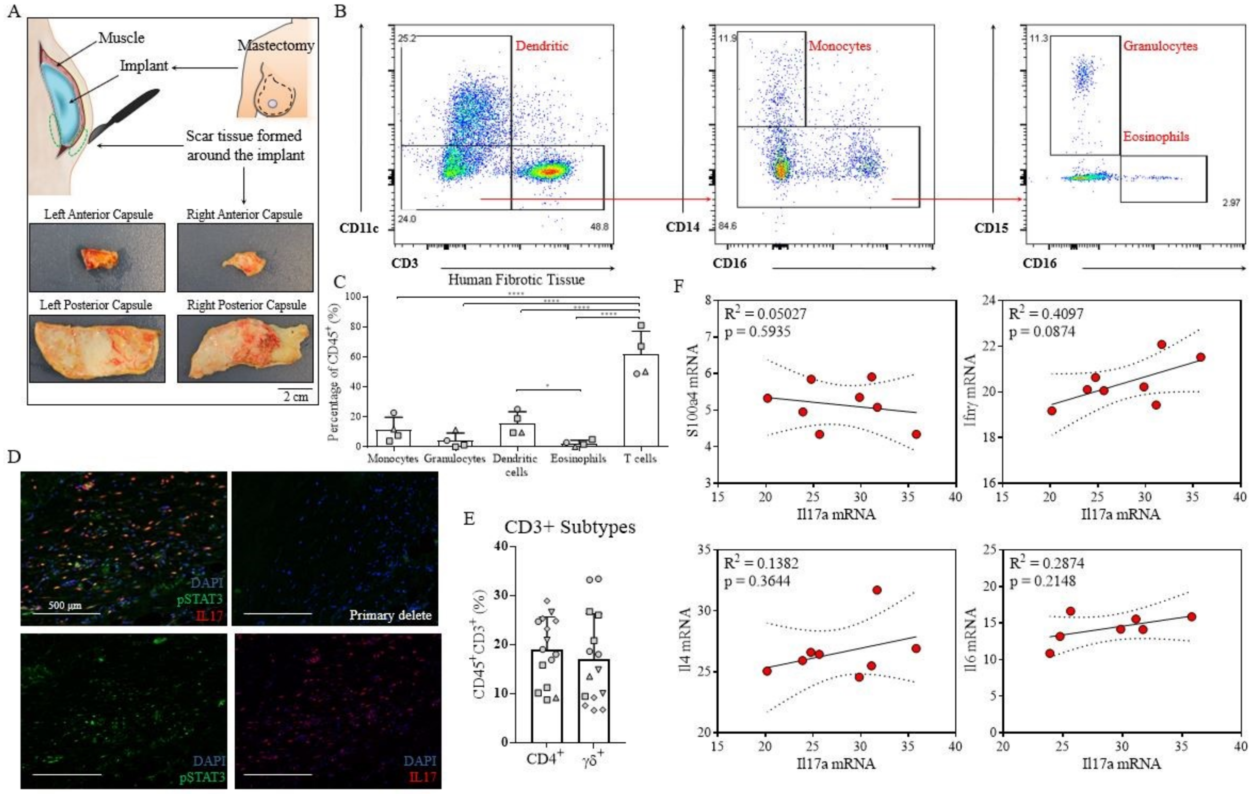


**Figure S1.** (**A**) Illustration of human fibrotic capsule extraction and gross images of the implants (**B, C**) Representative flow cytometry plot and quantification of myeloid derived cells, including monocytes ($CD3^-CD11c^-CD14^+CD16^-$), granulocytes ($CD3^-CD11c^-CD14^+CD16^+CD15^+$), eosinophils ($CD3^-CD11c^-CD14^+CD16^+CD15^-$), dendritic cells ($CD3^-CD11c^+$), and lymphoid derived T cells ($CD3^+CD11c^-$). The number of T cells was significantly higher than the number of myeloid cells in 3 patients. (**D**) Immunofluorescence imaging showing pSTAT3 and IL17 in human fibrous capsule, followed by single staining controls and primary delete. (**E**) Quantification of the percentage of $CD4^+$ T cells and $\gamma\delta^+$ T cells in 5 patients. (**F**) Correlation of qRT-PCR gene expression between Il17a mRNA and fibrosis-associated genes, including S100a4, Ifnγ, Il4, and Il6. Data are means ± SD, n = 3 (C), n = 8 (D), n = 5 (F) ANOVA [(C) (D) and (F)]: ****P < 0.0001, ***P < 0.001, **P < 0.01, *P < 0.05.

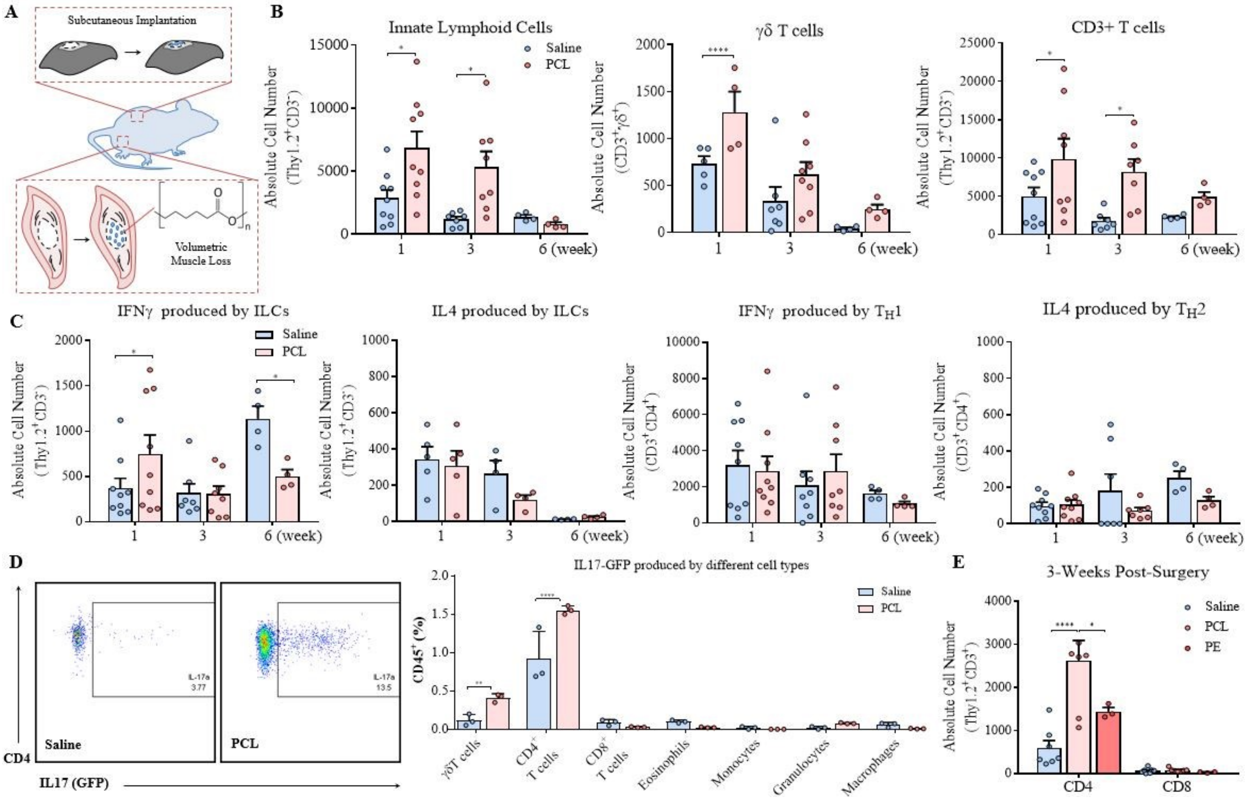


**Figure S2.** (**A**) Illustration of volumetric muscle loss (VML) and subcutaneous (SQ) implants in the murine model. (**B**) Flow cytometry analysis of the total number of ILCs, $\gamma\delta^+$ T cells, and $CD3^+$ T cells at 1, 3, and 6-weeks post-surgery. (**C**) Kinetics of IFNγ and IL4 expression by ILCs, $\gamma\delta^+$ T cells, and $CD4^+$ T helper cells at various post-injury time points. (**D**) Quantification of IL17A expression across different cell types 3-weeks post-surgery in an IL17A-GFP reporter mice. (**E**) Comparison of $CD4^+$ and $CD8^+$ T cells at 3-weeks post-injury with PCL and PE implant. Data are means ± SD, ANOVA [(B), (C) and (D)]: ****P < 0.0001, ***P < 0.001, **P < 0.01, *P < 0.05.

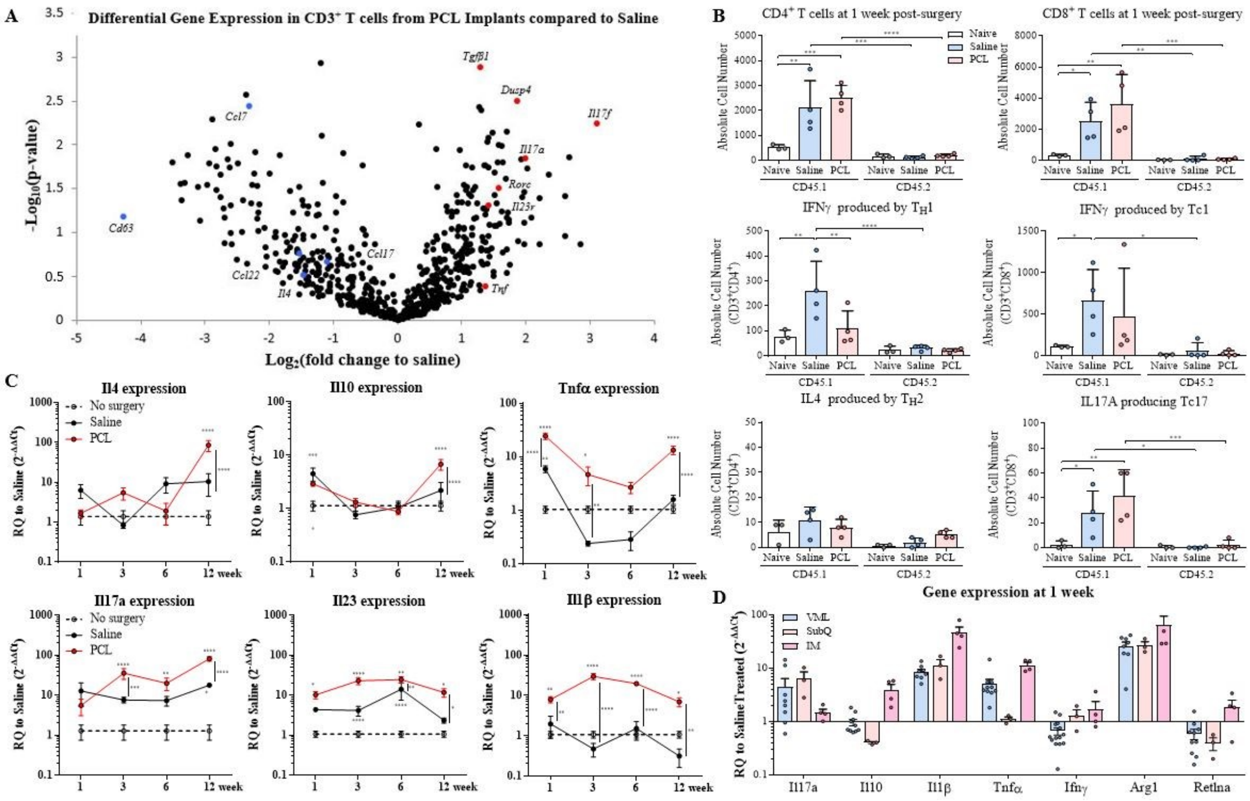


**Figure S3.** (**A**) $CD3^+$ T cells were sorted for gene expression analysis using the NanoString platform. Volcano plot of genes differentially regulated in PCL-derived T cells compared to saline (no implant) control at day 7 post-surgery. Type 17-associated gene expression differences are highlighted. (**B**) Total number of CD4 and CD8 infiltration, $T_H1$, and $T_H2$, Tc1, Tc17 from CD45.1 (WT donor) and CD45.2 (OTII-$Rag^{-/-}$ donor) bone marrow chimera mice 1 week after VML. (**C**) qRT-PCR gene expression normalized to healthy muscle control showing kinetics of Il4, Il10, Il17a, Il23, Tnfα, and Il1β transcripts. (**D**) Gene expression analysis comparing levels of different cytokines between the VML model and subcutaneous (SQ) implant, and intramuscular injection (IM) model at 1-week post-injury. Data are means ± SEM, ANOVA [(B) and (C)]: ****P < 0.0001, ***P < 0.001, **P < 0.01, *P < 0.05.

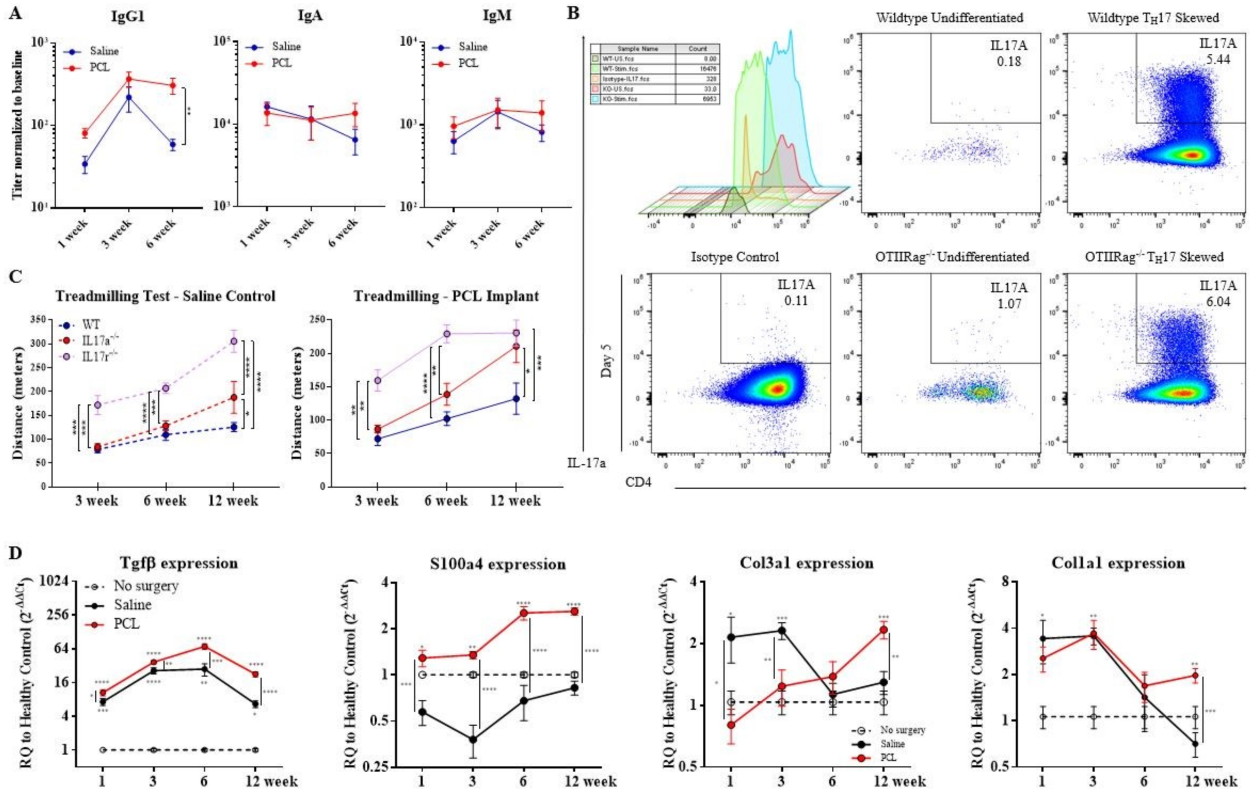


**Figure S4.** (**A**) IgG1, IgM, and IgA antibody titers were detected using an enzyme-linked immunosorbent assay (ELISA) system on serum collected 1, 3, and 6-weeks post-surgery and implantation. (**B**) *In vitro* differentiation of $T_H17$ cells of OTII-Rag$^{-/-}$ compared to WT naïve CD4$^{+}$ cells. (**C**) Treadmill exhaustion assay 3, 6, and 12-weeks post-implantation of WT, IL17A$^{-/-}$, and IL17RA$^{-/-}$ mice. (**D**) qRT-PCR gene expression normalized to healthy muscle controls showing kinetics of Tgfβ, S100a4, type I and III collagen progression in PCL implant mice compared to saline control overtime. Data are means ± SEM, ANOVA [(A), (C) and (D)]: ****P < 0.0001, ***P < 0.001, **P < 0.01, *P < 0.05.

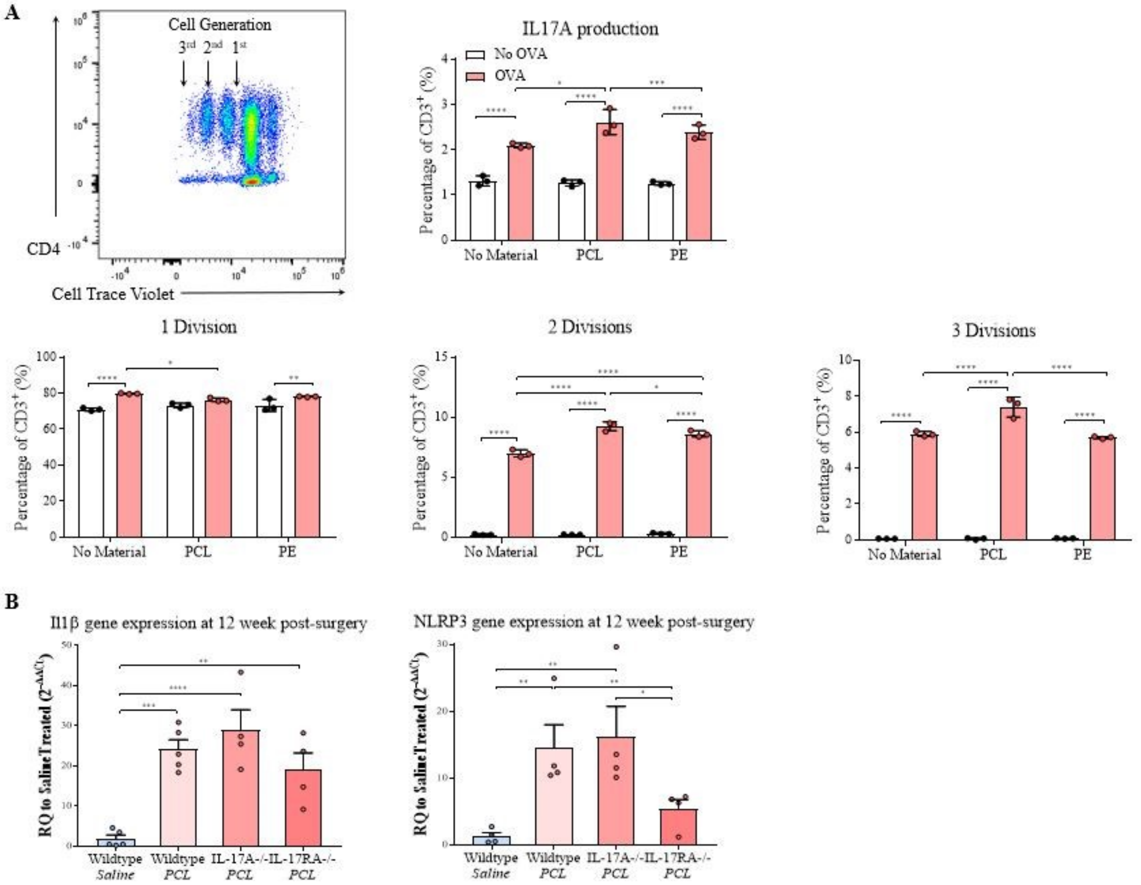


**Figure S5.** (**A**) Proliferation assay of T cells co-cultured with spleenocytes from OT-II transgenic mice cultured with different materials (PCL or PE) and OVA antigen. Representative plots for proliferation assay were shown. In addition, level of IL-17A were also quantified via intracellular staining. (**B**) qRT-PCR gene analysis of fibrotic markers including Il1β and Nlrp3 in WT, IL17A$^{-/-}$, and IL17RA$^{-/-}$ mice at 12-weeks post-surgery. Data are means ± SD, n = 3 [A], n = 4 [B], ANOVA (A, B): ****P < 0.0001, ***P < 0.001, **P < 0.01, *P < 0.05.

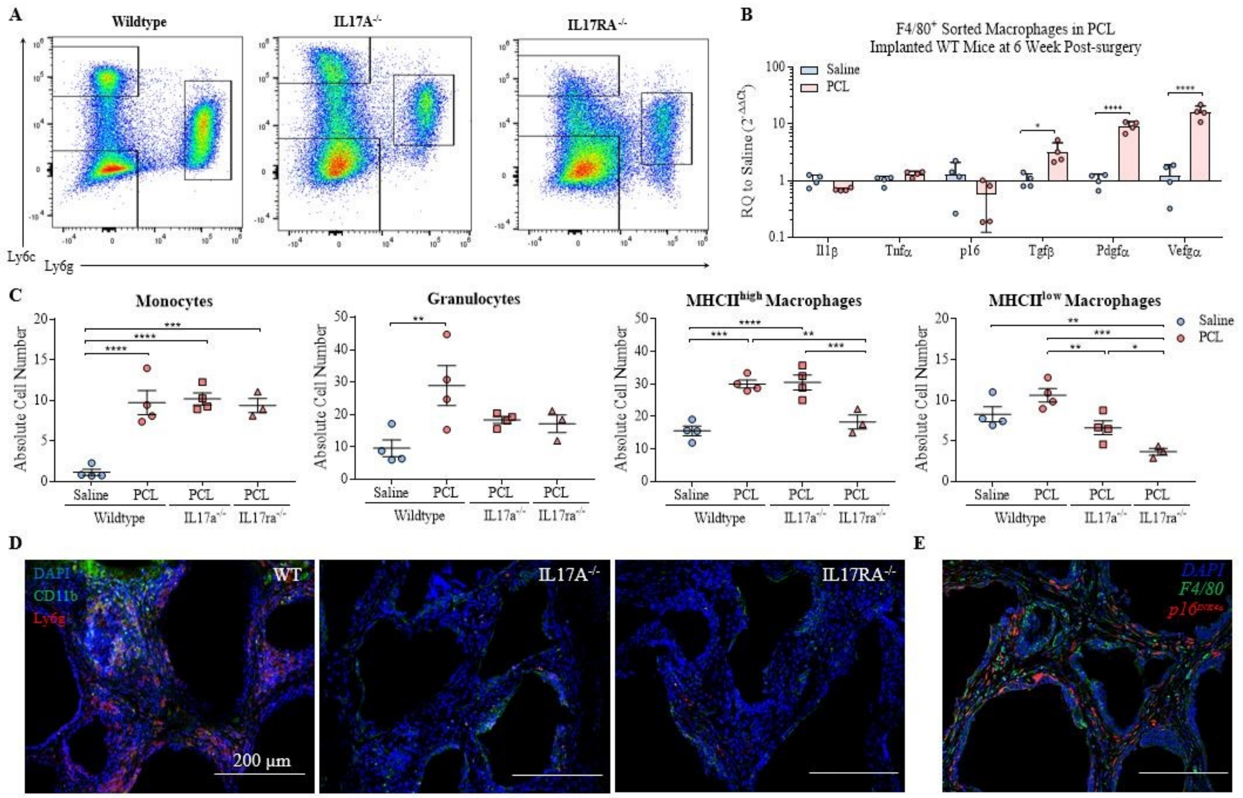


**Figure S6.** (**A**) Flow cytometry representative plots comparing different populations of myeloid cells in WT, IL17A$^{-/-}$, and IL17RA$^{-/-}$ mice. (**B**) Gene expression analysis of F4/80$^{+}$ cells sorted from PCL implanted WT mice 6-weeks post-surgery. (**C**) Quantification of granulocytes (CD11b$^{+}$Ly6intLy6g$^{high}$), MHCII$^{high}$ macrophages (CD11b$^{+}$Ly6c$^{low}$Ly6g$^{low}$MHCII$^{high}$F4/80$^{+}$), MHCII$^{low}$ macrophages (CD11b$^{+}$Ly6c$^{low}$Ly6g$^{low}$MHCII$^{low}$F4/80$^{+}$), and monocytes (CD11b$^{+}$Ly6c$^{high}$Ly6g$^{low}$) in WT, IL17A$^{-/-}$, and IL17RA$^{-/-}$ mice after 3 weeks with PCL implants. (**D**) Immunofluorescence imaging of CD11b and Ly6g around PCL implants in WT and IL17 signaling deficient mice 12-weeks post-surgery. (**E**) Immunofluorescence imaging of F4/80 and p16$^{INK4a}$ in WT mice 12-weeks post-surgery. Data are means ± SEM, n = 4, ANOVA (B): ****P < 0.0001, ***P < 0.001, **P < 0.01, *P < 0.05.

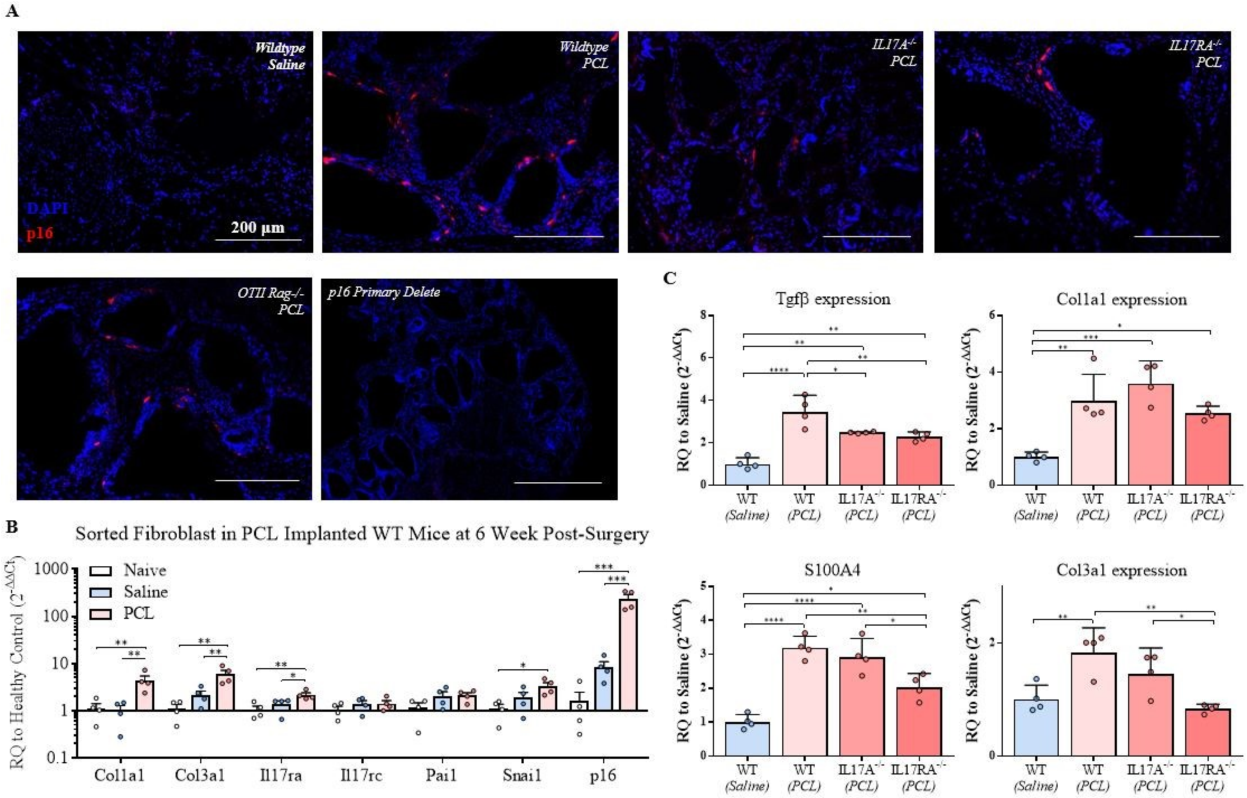


**Figure S7.** (**A**) Immunofluorescence staining of p16$^{INK4a}$ (red) in WT, IL17A$^{-/-}$, and IL17RA$^{-/-}$, OTII-Rag$^{-/-}$ mice at 6-weeks post-injury, followed by primary antibody delete control. (**B**) Gene expression analysis of fibroblasts sorted from PCL implanted WT mice 6-weeks post-surgery. (**C**) qRT-PCR gene analysis of fibrotic markers in the whole tissue including Tgfβ, S100a4, and type III collagen in WT, IL17A$^{-/-}$, and IL17RA$^{-/-}$ mice at 12-weeks post-surgery. Data are means ± SEM, n = 4, ANOVA (B, C): ****P < 0.0001, ***P < 0.001, **P < 0.01, *P < 0.05.

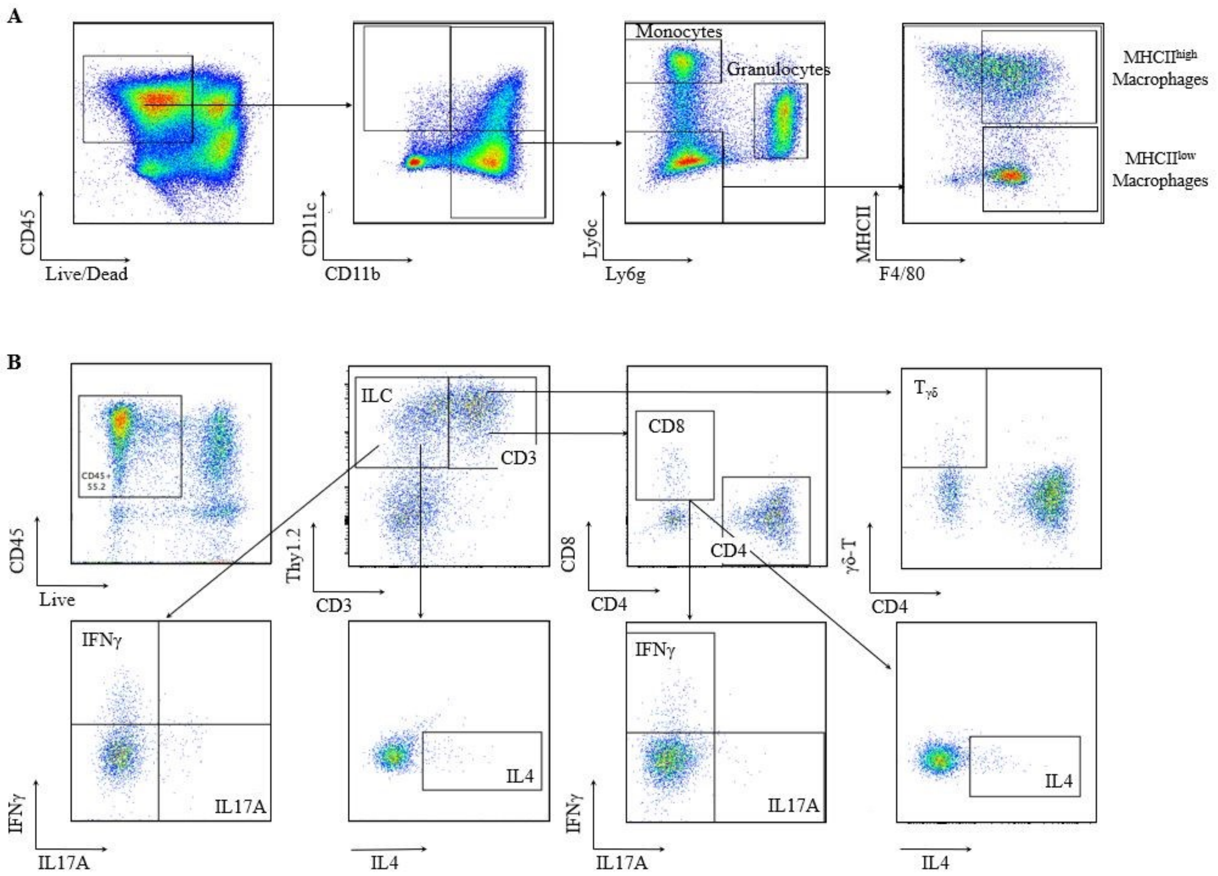


**Figure S8.** Flow cytometry gating strategies for myeloid (**A**) and lymphoid (**B**) populations.
